# Limits on visual awareness of object targets in the context of other object category masks: Investigating bottlenecks in the continuous flash suppression paradigm with hand and tool stimuli

**DOI:** 10.1101/363515

**Authors:** Regine Zopf, Stefan R. Schweinberger, Anina N. Rich

## Abstract

The continuous flash suppression (CFS) task can be used to investigate what limits our capacity to become aware of visual stimuli. In this task, a stream of rapidly changing mask images to one eye initially suppresses awareness for a static target image presented to the other eye. Several factors may determine the breakthrough time from mask suppression, one of which is the overlap in representation of the target/mask categories in higher visual cortex. This hypothesis is based on certain object categories (e.g., faces) being more effective in blocking awareness of other categories (e.g., buildings) than other combinations (e.g., cars/chairs). Previous work found mask effectiveness to be correlated with category-pair high-level representational similarity. As the cortical representations of hands and tools overlap, these categories are ideal to test this further, as well as to examine alternative explanations. For our CFS experiments, we predicted longer breakthrough times for hands/tools compared to other pairs, due to the reported cortical overlap. In contrast, across three experiments, participants were generally faster at detecting targets masked by hands or tools compared to other mask categories. Exploring low-level explanations, we found that the category average for edges (e.g., hands have less detail compared to cars) was the best predictor for the data. This low-level bottleneck could not completely account for the specific category patterns and the hand/tool effects, suggesting there are several levels at which object category-specific limits occur. Given these findings, it is important that low-level bottlenecks for visual awareness are considered when testing higher-level hypotheses.

## Introduction

We live in a rich and complex world and are typically surrounded by many different types of objects. However, our ability to perceive and become aware of visual stimuli is limited (Cohen, Dennett, & Kanwisher, 2016; C. Y. Kim & Blake, 2005; Treisman & Gelade, 1980; Wolfe, Cave, & Franzel, 1989). The mechanisms that underlie these limitations have typically been associated with perceptual inhibitory interactions in low-level visual cortex (Tong, Meng, & Blake, 2006) or with limited attentional capacities in the fronto-parietal network (Desimone & Duncan, 1995). Understanding how our capacity limits are generated is a key question in understanding awareness.

The continuous flash suppression (CFS) task can be used to investigate what limits our capacity to become aware of visual stimuli. This paradigm has been used by several researchers to study processing across different visual object types (Almeida, Mahon, Nakayama, & Caramazza, 2008; Cohen, Nakayama, Konkle, Stantic, & Alvarez, 2015; Hesselmann, Darcy, Ludwig, & Sterzer, 2016; Ludwig, Kathmann, Sterzer, & Hesselmann, 2015; Stein, Sterzer, & Peelen, 2012). The CFS task involves presenting a stream of rapidly changing mask images to one eye and a static target image to the other eye (Tsuchiya & Koch, 2005; Tsuchiya, Koch, Gilroy, & Blake, 2006). The separate images are fused in the brain and participants tend to experience a continuous stream of the flashing mask images. This initially suppresses awareness for the static target image, but this may become visible after some time.

Several factors may determine the time it takes targets to break through mask suppression. One proposed factor is the categorical similarity in higher visual cortex. This hypothesis is based on the finding that in the CFS task certain object categories (e.g., faces) are more effective in blocking awareness of other categories (e.g., buildings) than other combinations (e.g., cars/chairs), which was found to correlate with category-pair representational similarity in higher visual cortex (Cohen et al., 2015). Typically, the mask images in CFS tasks are Mondrian-style patterns because these are highly effective in target suppression. Cohen et al. (2015), however, used object stimuli as masks, which allows a nice measure of how long it takes targets to break through the visual suppression for different types of category combinations (e.g., a building target breaking through suppression from masking faces). This approach can therefore be used to explore the factors that determine breakthrough times in CFS.

Cohen and colleagues conducted a series of experiments using a range of paradigms that involve the presentation of multiple object stimuli either spatially separated (visual search paradigm: Cohen, Alvarez, Nakayama, & Konkle, 2017; visual memory paradigm: Cohen, Konkle, Rhee, Nakayama, & Alvarez, 2014) or overlapping (visual masking and continuous flash suppression paradigms: Cohen et al., 2015). In each experiment, they found that certain category combinations (e.g., faces/buildings) were more effective in interfering with target processing than others (e.g., cars/chairs). Reduced performance for specific category pairs was in each case predicted by more representational similarity in higher visual cortex for the paired objects (Cohen et al., 2014, 2015, 2017). Thus, they made a general proposal that the extent to which high-level visual categories (such as faces, bodies and chairs) have spatially separable neural representations predicts the capacity to simultaneously process multiple visual stimuli. This high-level representational architecture theory thus offers one mechanism for our limitations on processing object information in the context of competing visual input. This may therefore be one contributor to the limits on visual awareness in the CFS paradigm.

As hand and tool representations in higher visual cortex are close to each other and partly overlap (Bracci, Cavina-Pratesi, Ietswaart, Caramazza, & Peelen, 2012), these categories provide ideal stimuli to test the idea that neural overlap might contribute to CFS breakthrough times. On the basis of the high-level representational architecture model, we can make specific predictions about the behavioural task performance for paradigms involving hands and tools stimuli. Although Cohen et al. (2015) quantified neural similarity in a separate fMRI experiment, here, we used existing neural data showing the overlap of hands and tools to predict breakthrough times for hand/tool pairs compared to other pairs (e.g. hands/cars). Further, previous representational similarity analyses have shown that neural representation of hands is somewhat closer to that for small than large objects (Kriegeskorte et al., 2008). Thus, the high-level representational architecture model also predicts reduced task performance for hands and small objects compared to hand and large object pairs.

Hands are commonly involved in our interactions with the external world. To be successful in these interactions we have to be able to quickly and dynamically update our representations of target objects in the context of competing visual input. Each interaction involves at least two objects - at least one effector (e.g. a hand or a hammer) and one target (e.g. a phone or a nail). Thus, finding out more about the visual perceptual limitations for processing of objects in the context of bodily information is also important for understanding perceptual factors that play a role in interactions between body and environment.

One of the crucial aspects of comparing category-level interactions is to select stimuli that have similar variability. Our hand category consisted of just one type of stimulus (basic category level) with many different exemplars of hands (Rosch, Mervis, Gray, Johnson, & Boyes-Braem, 1976). In other higher-level category sets of tools, small objects and large objects (superordinate categories), there is a much greater variability between category exemplars (e.g., a tool could be a hammer, pliers, screwdriver etc). We therefore also included other basic category sets that consisted of only one type of object (hammers, phones and cars) to provide an appropriate control for category-level and variance in the hand condition.

We tested if the high-level neural representational architecture model can predict CFS breakthrough time rank-orders. Following Cohen et al. we tested pairs of stimuli in the CFS paradigm. Here, we use ‘hands/tools’ to indicate a hand target competing with tool masks as well as a tool target competing with hand masks. Neural hand and tool representations overlap (Bracci et al., 2012) and neural representations for small objects are further away from hand and tool representations then they are from each other; large object representations are even further away in neural representational similarity space (Kriegeskorte et al., 2008). Thus, we derived and tested the following simple models of rank-order predictions (also see Figure 2): *1^st^ Rank*: The fastest breakthrough times for hand/large object and tool/large object pairs. *2^nd^ Rank*: Somewhat slower breakthrough times for hand/small objects, tool/small object and small object/large object pairs because these representations are closer together. *3^rd^ Rank*: Even slower breakthrough times for hand/tool pairs due to the overlap and *4^th^ Rank*: slowest breakthrough times for within-category pairs because the high-level neural architecture model predicts the most overlap for stimuli from the same category.

In addition to predictions based on neural representation, we can also test the contribution of other factors to CFS breakthrough times using this paradigm. Here, again, hands provide useful stimuli. Although we globally matched (i.e., for the entire image) characteristics such as size, colour, contrast, luminance and spatial frequency content across all images, there are still category-specific local image characteristics that could influence breakthrough times. For example, hands, due to their shape, might not cover the entire target area, which could facilitate target breakthrough from suppression. We calculated the average percentage of object across the target area and also across the entire image for each mask category, allowing us to create object-coverage models as predictors for breakthrough times for different category masks.

A further image characteristic that could influence breakthrough times is the amount of edges (local clustering of contrast changes). In fact, it has previously been suggested that the amount of edges could determine CFS because increased spatial density of Mondrian masks result in longer breakthrough times (Drewes, Zhu, & Melcher, 2018). However, in this previous study, their manipulation of spatial density changed both edge content and global spatial frequency content, contrast and luminance. Thus, it was not possible to disentangle if increased CFS breakthrough times were due to increased edge content or due to increased contrast energy. Here we used images with matched global spatial frequency content, contrast and luminance, which allowed us to specifically test the effect of edge content.

In summary, we used CFS to test the contribution of different factors, ranging from high-level neural representation to low-level object coverage and edge content models, to breakthrough times for different object category pairs. The results have implications for our understanding of visual awareness and multiple object perception.

## Methods Experiment 1

In Experiment 1 we tested if known characteristics of neural high-level object representations predict CFS breakthrough times for the categories of: hands, tools, hammers, small objects, phones, large objects and cars.

### Participants

As in Cohen et al. (2015), we tested 20 participants. Our sample consisted of 17 females and 3 males (M = 21.05 years, SD=2.65, range 18-27 years, 2 left-handed according to self-report, 12 right-dominant eye). All participants had normal or corrected-to-normal vision (contact lenses only) and received $20 for their participation. The data of one participant had to be replaced due to technical difficulties that resulted in an incomplete data set. This research was conducted in accordance with the ethical standards laid down in the 1964 Declaration of Helsinki and was approved by the Macquarie University Ethics Review Committee (Human Research). Written informed consent was obtained from all participants prior to the start of the experiment.

### Stimuli

We prepared our stimuli in line with the steps in Cohen et al. (2015) to minimize low-level differences between stimulus sets. We selected visually variable stimuli (see Figure 1A and 1B, the complete set of stimuli is available here: https://bit.ly/2Krq7Ab) and, also in the interest of high stimulus set variability, selected 60 stimuli per category (Cohen et al. 2015 used 30 stimuli) and presented these in both mirrored and non-mirrored versions. Stimuli for all three experiments were prepared at the same time and selected from our own databases as well as from the Bank of Standardized Stimuli (BOSS, sites.google.com/site/bosstimuli/, Brodeur, Dionne-Dostie, Montreuil, & Lepage, 2010), Konklab image sets (http://konklab.fas.harvard.edu/#, Konkle & Oliva, 2012) and the Tarrlab (www.tarrlab.org, Stimulus images courtesy of Michael J. Tarr, Center for the Neural Basis of Cognition and Department of Psychology, Carnegie Mellon University. Funding provided by NSF award 0339122). All stimuli were centred and gray-scaled. We normalized the intensity histogram (i.e., contrast and luminance) across all images and power at all spatial frequencies and orientations using the SHINE toolbox (Willenbockel et al., 2010).

**Figure 1.**
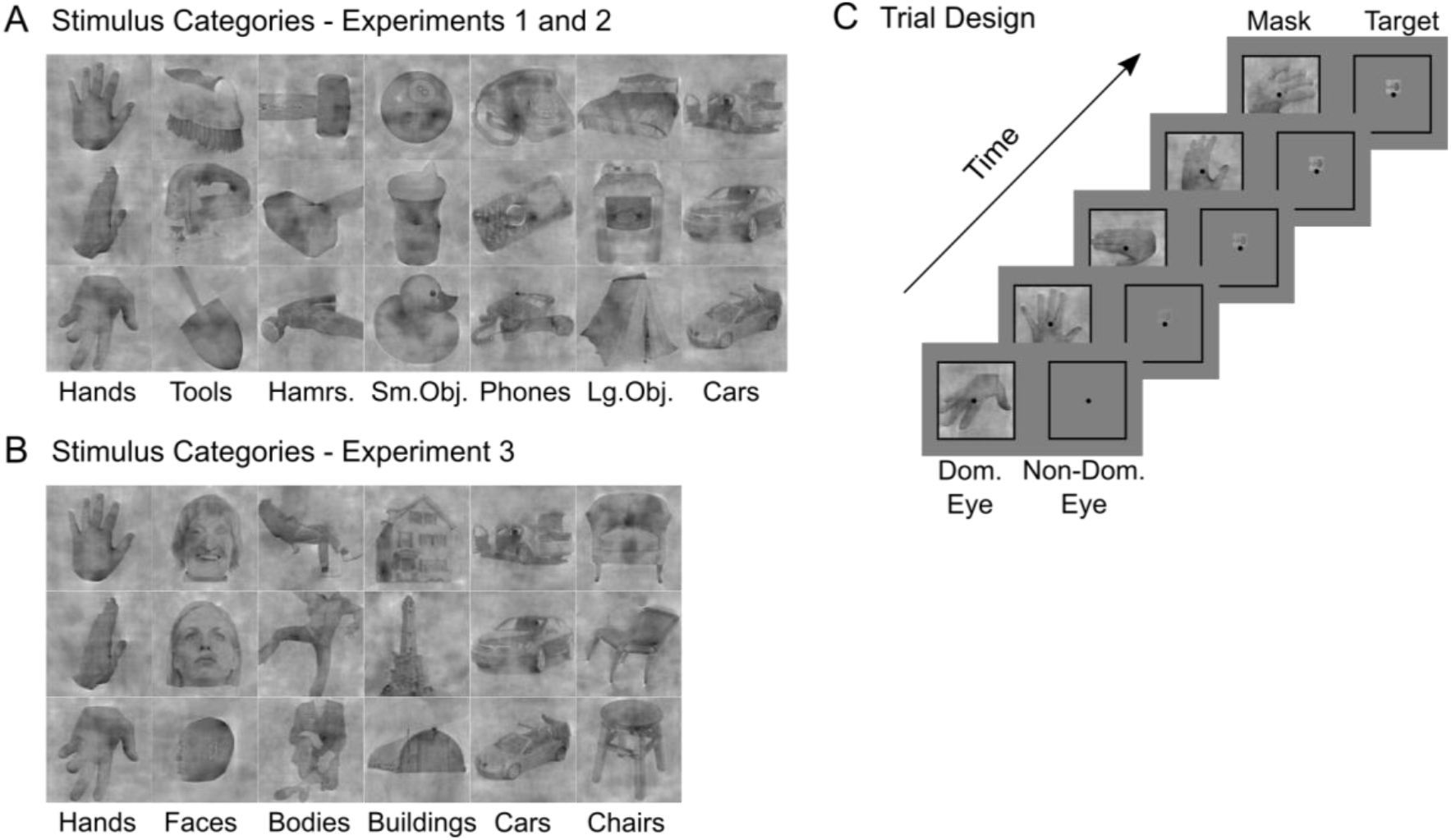
Stimuli and Continuous Flash Suppression (CFS) trial design. A) In Experiments 1 and 2, we presented hands, tools, hammers, small objects, phones, large objects and cars. Three stimulus exemplars for each set are depicted. B) In Experiment 3, we presented hands, faces, bodies, houses, cars and chairs. Three stimulus exemplars for each set are depicted. C) On each CFS task trial, mask images were presented in the dominant eye and target images in the non-dominant eye.

### Continuous Flash Suppression Task

Using the CSF task, we measured the time for target categories to break through suppression from distracting mask categories. We determined eye dominance for each participant before the experiment using the Miles test (Miles, 1930). We used a mirror stereoscope (ScreenScope, Stereoaids, Albany, Australia, stereoaids.com.au) to present mask stimuli to the dominant eye and target stimuli to the non-dominant eye (Figure 1C). Before running the task, we presented a frame and fixation dot to each eye and asked participants if these were clearly visible with each eye and if they overlapped when both eyes were open. We adjusted the stereoscope mirror angles as needed. The mask images were large (16.5° visual angle) and centred in the middle of the screen. The target images were small (3° visual angle) and presented in the centre of the screen either just above or below a small central fixation dot. To facilitate fusion of target and mask images, the target image was presented through a square Gaussian aperture (i.e., the outer 5% of the image was smoothed) and a thin black frame with the size of the mask image was presented to both eyes. This differed slightly from Cohen et al. (2015), who used a circular aperture, to ensure that we did not cut off any part of the target stimuli (particularly for the hands). A new mask image was shown every ~167 ms (6 Hz). This has been shown to be the optimal temporal frequency for suppression in other CFS studies (Drewes et al., 2018; Zhu, Drewes, & Melcher, 2016), but is actually a slower rate than Cohen et al. (2015), who used ~117 ms (8.5 Hz); nevertheless we found comparable breakthrough times (see Experiment 3). A trial started with the presentation of the black fixation dot for 250 ms which then turned red for 200 ms to alert participants that the trial was about to start. From the start of the trial, the static target gradually became more visible (0% to 100% opacity in 13 steps with each new mask presentation) over ~2170 ms. After that, the mask gradually became less visible (100% to 40% opacity in 37 steps with each new mask presentation) over ~6180 ms. If there was no response, the trial ended after ~8350 ms. Over the course of the trial, up to 50 different mask images were randomly selected from the set of 120 options (mirrored and non-mirrored versions of our 60 unique stimuli per category).

Participants were instructed to detect the appearance of the small target item and respond with the space bar when they detected it. This stopped the stimulus presentation, and they then gave a forced choice response indicating if the target was presented above or below fixation by pressing one of two keyboard keys (‘1’ or ‘2’). Participants were instructed to give the first response as fast as possible and the second response as accurately as possible. Participants completed 10 practice trials before each block and received visual feedback (‘correct’, ‘incorrect’ or ‘no response’) for 1 s after every trial. The next trial started immediately after the feedback screen.

Mask stimuli were presented in blocks – separately for each of the seven category sets (hands, tools, hammers, small objects, phones, large objects, cars). For 10 participants we used a random order of the 7 conditions, and for the other 10 participants, we used the reverse of those 10 orders. In each mask block, there were 20 target trials for each category; in contrast to Cohen et al. 2015, we also presented targets from the same category as the mask. For each category, in half of the trials (10 target trials) the target appeared above and in the other half below fixation. In total, there were 140 trials within each block; target categories and target locations presented in a random order. There were two breaks within each mask block and a break between mask blocks. The experiment took approximately 75 minutes to complete. Stimulus presentation was controlled with MATLAB (The MathWorks, Natick, MA) and the Psychtoolbox (Brainard, 1997; Pelli, 1997). Stimuli were presented on an ASUS monitor with a refresh rate of 60 Hz and a screen resolution of 1920 x 1080. The eye to monitor distance was approximately 57 cm.

### Data Analysis

Our variable of interest was the breakthrough reaction time, which is how long it took participants to press the space bar to indicate target detection. Trials without a response were excluded from analysis (0.30% of trials). We then also excluded trials on which the subsequent location response was incorrect: participants were 98.60% (SD=1.07) correct across all trials. Finally, trials with response times < 300 msec or > three standard deviations from the participant’s mean across all trials were also excluded, leading to the removal of a further 1.64% of the trials.

We developed a novel version of analysis methods that are frequently used in the context of neuroimaging data representational similarity analyses (RSA, Nili et al., 2014). First, we used 2D mosaic plots for visualisation of the breakthrough reaction times (see Figures 2–5). This allows display of the matrix of breakthrough reaction times for all mask and target combinations in one plot and facilitates examination of the data pattern. Second, we employed correlation analyses to assess the similarity between specific models and the data. This allowed us to assess and compare the relatedness of different candidate models. Because our model predictions did not specify linear relationships but instead simply a rank order for category pairs (e.g., slower breakthrough times for hand/tool pairs compared to hand/large object pairs), we used rank-order correlations. Specifically, we used Kendall’s rank correlation coefficient τ_A_ (‘tau-a’), which is the proportion of values that are consistently ordered in both variables and is suitable for comparisons of models that predict tied ranks (Nili et al., 2014). We programmed and ran these analyses in MATLAB. In addition to the paired data-model correlation tests, we also ran partial correlation analyses to test data-model correlations while controlling for other variables. To run partial nonparametric (Kendall’s rank) correlation analyses we used the R package ppcor (S. Kim, 2015). In addition, as a sanity check, we also ran these correlation and partial-correlation analyses using the available MATLAB functions for nonparamatric (Spearman’s rank) correlations and found somewhat higher correlation coefficients but overall a consistent pattern of results.

**Figure 2.**
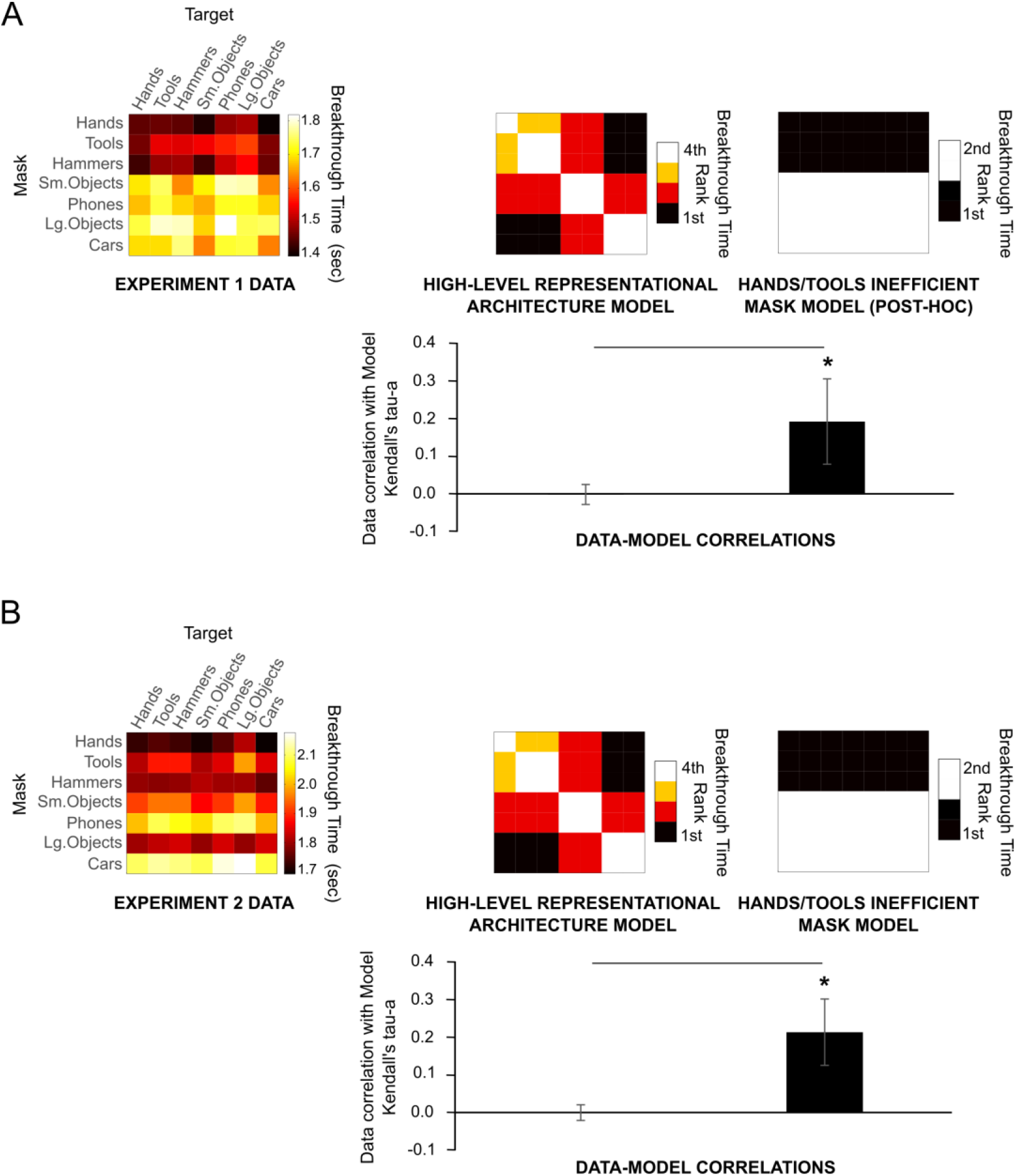
Results for high-level category-specific models Experiment 1 (A) and Experiment 2 (B). Mosaic data plots depict breakthrough times for all mask and target combinations. Darker colours are faster and lighter colours slower breakthrough times. Across both experiments fast breakthrough times are found for targets when hands were masks. Mosaic model plots depict the predictions for the rank order of mask and target combinations (e.g., hands & cars fastest and hands & hands slowest breakthrough times). We found no significant data-model correlations for the high-level representational architecture model. We found significant correlations for the hand/tool inefficient mask model. Asterisks indicate significance of individual models and lines significant pairwise comparisons, both assessed using permutation tests. Multiple model comparison was accounted for using Bonferroni corrections. Error bars represent 95% confidence intervals.

**Figure 3.**
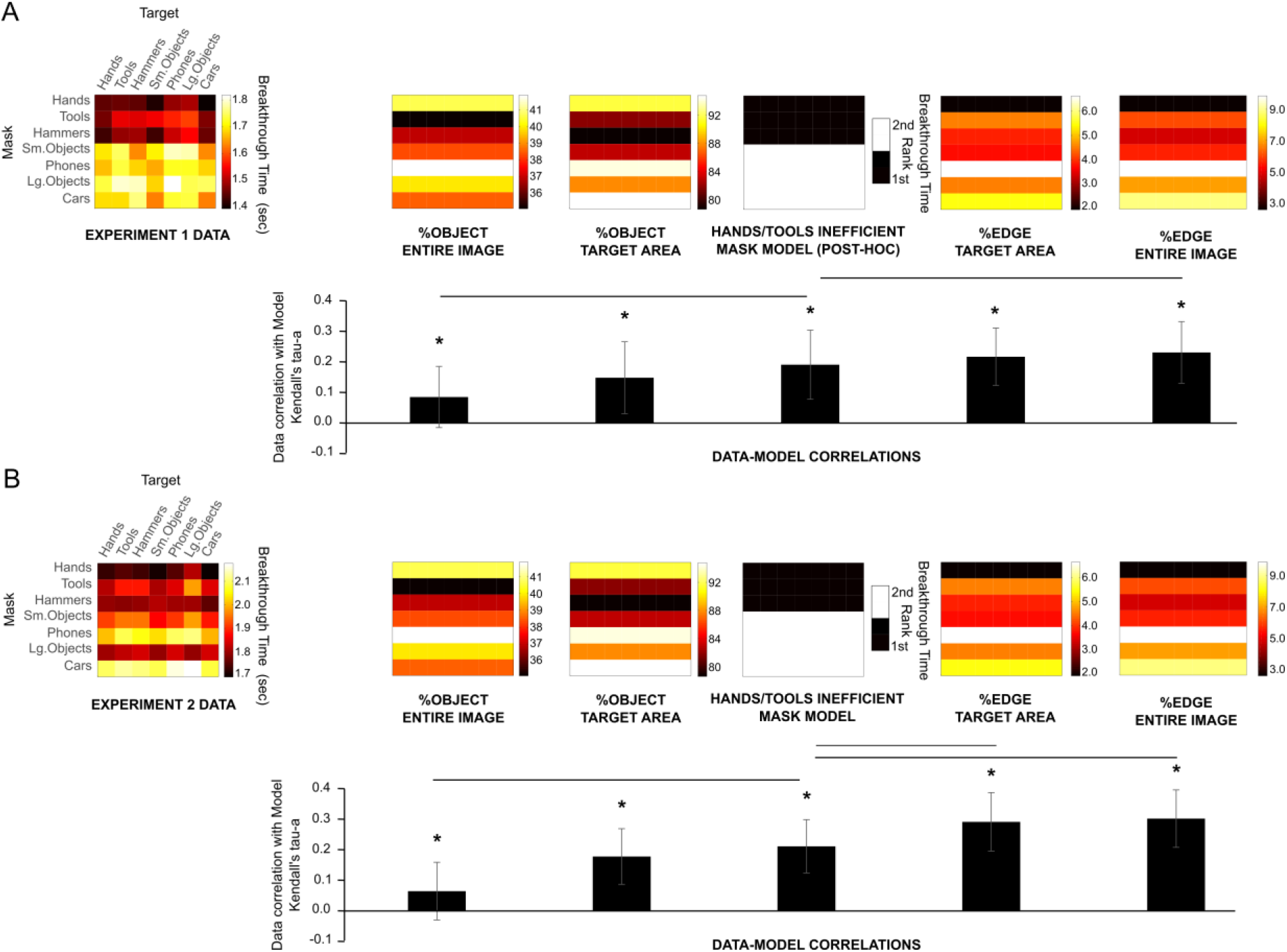
Results for local category-specific image characteristics for Experiments 1 (A) and 2 (B). We calculated category specific values for the amount of edges (local luminance changes) as well as the amount of image covered by the object both specifically in the target area and also for the entire image. For the amount of edges we found very similar or higher data-model correlations compared to the hands/tools inefficient mask model. Asterisks indicate significance of individual models and lines significant pairwise comparisons, both assessed using permutation tests. Multiple model comparison was accounted for using Bonferroni corrections. Error bars represent 95% confidence intervals.

**Figure 4.**
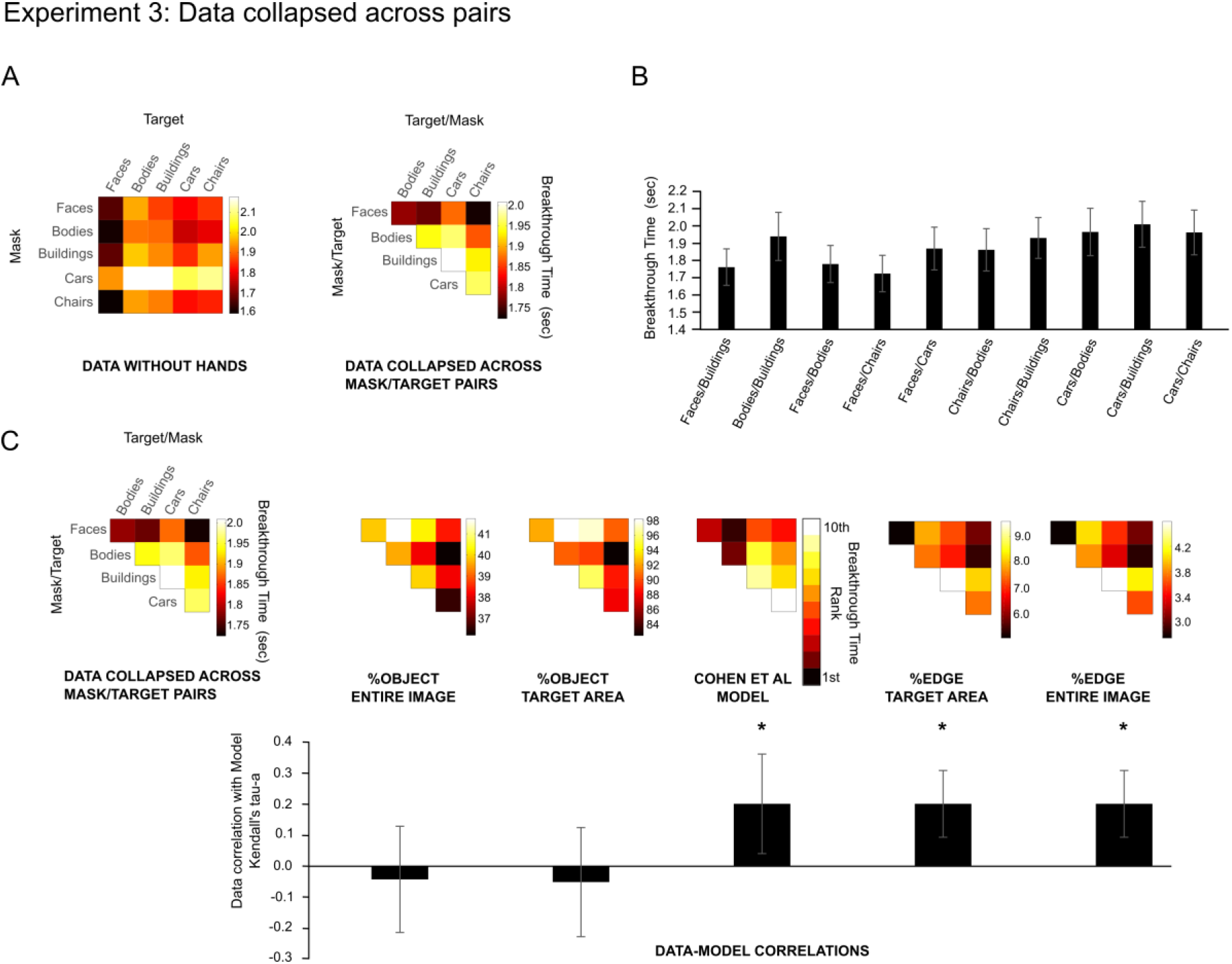
Data and models for Experiment 3 as in Cohen et al. (2015). A) Only includes categories of faces, bodies, buildings, cars and chairs; within-category mask and target pairs are excluded and we collapsed data across mask and target pairs (e.g., face mask and body target averaged with body mask and face target) as in Cohen et al. (2015). B) The collapsed mask and target pairs as ordered in Cohen et al. (2015). We found a significant correlation with the rank-order Cohen et al. (2015) reported previously (Spearman’s rho = 0.71, p=0.019). C) Cohen et al. (2015) order model (which previously showed high correlations with the high-level neural category representations), edge-content and object-coverage models as well as data-model correlations. The Cohen et al. (2015) model and edge content models had the highest data-model correlation coefficients and were not significant different. Asterisks indicate significance of individual models assessed using permutation tests. Multiple model comparison was accounted for using Bonferroni corrections. Error bars represent 95% confidence intervals.

**Figure 5.**
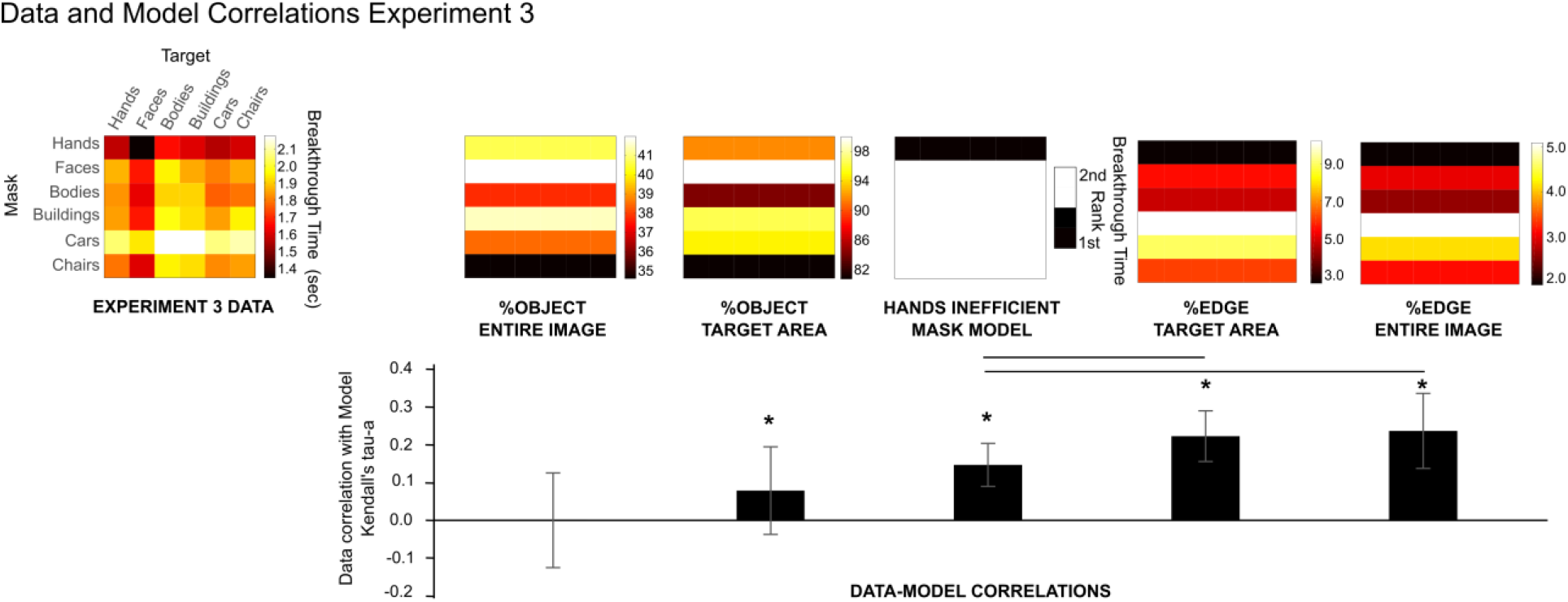
Data, models and correlation analyses for Experiment 3. Models include the hand inefficient mask model and models for category specific edge content as well as the amount of image covered by the object both specifically in the target area and also for the entire image. The edge-content models showed the highest data-model correlations of all the models. Asterisks indicate significance of individual models and lines significant pairwise comparisons, both assessed using permutation tests. Multiple model comparison was accounted for using Bonferroni corrections. Error bars represent 95% confidence intervals.

We used permutation tests for statistical inference. To estimate the sampling distribution of the correlation coefficients under the null hypothesis, we randomly shuffled trial labels for each participant’s data set of target/mask pair breakthrough times and then calculated condition means and data-model correlation coefficients for 10,000 times. We calculated two-sided p-values as the proportion of absolute sampled correlation coefficients that were greater or equal to the absolute observed correlation coefficient. Exact p-values were calculated with the R function permp (statmod package) (Phipson & Smyth, 2010). We corrected our analyses for multiple comparisons using Bonferroni adjusted alpha significance levels.

## Results Experiment 1

The breakthrough time data pattern for all mask and target combinations did not resemble the predicted high-level representational architecture model (Figure 2A). Hand and tool combinations did not lead to slower breakthrough times compared to other hand and object categories. In contrast, they resulted in some of the fastest breakthrough times (dark colours). Overall, we did not find significant data-model correlations for the high-level representational architecture model (tau-a mean=−0.002, *p*=0.93).

What can be clearly seen is that there are general mask effects: hand and tool masks resulted in the fastest mean breakthrough time across different target combinations (hands mean RT=1447 ms, SD=434 ms; tools mean RT=1526 ms, SD=508 ms; hammers mean RT=1475 ms, SD=488 ms; small objects mean RT=1712, SD=517 ms; phones mean RT=1694, SD=450 ms; large objects mean RT=1759, SD=782 ms; cars mean RT=1695, SD=510 ms). This suggests that hands and tools both are relatively inefficient masks. This is not just due to the category level (basic vs supraordinate) as it is present for both levels (e.g., the tool category as well as the hammer) compared to all other categories. If we test a post-hoc model that has hands and tools as relative inefficient masks, there is a significant data-model correlation (exploratory analysis; Bonferroni corrected significance threshold of *p*=0.025, tau-a mean=0.191, *p*<0.001). These correlations were significantly higher than the correlations for the high-level representational architecture model (permutation test p-values for the difference: *p*<0.001).

These results show that target stimuli reach awareness faster when competing with hands or tools than when competing with other stimuli. We conducted a second experiment to further understand the mechanism for this effect and to test if we can replicate this pattern. It is possible that the effect is not due to differences in mask efficiency *per se*. Instead, it could be due to the strong association between hands or tools and controlling manual actions. Thus, viewing hands and tools over several seconds in the CFS paradigm could speed up manual motor breakthrough times (Longo & Haggard, 2009). This would then result in quicker reaction times to targets when these masks are used. In Experiment 2, we therefore used vocal (non-manual) responses to test if these hand and tool relative inefficient mask effects replicate.

## Experiment 2

In Experiment 2, participants gave vocal responses. We tested if the speed up of breakthrough reaction times in the context of hand and tool mask that we found in Experiment 1 replicates when using a non-manual response modality.

## Methods

We implemented the same methods as in Experiment 1 with the exceptions that we obtained vocal reaction times (see details below) and that we tested the predictions of both the high-level representational architecture and the hands/tools inefficient mask model (see Figure 2B).

### Participants

We tested 20 new participants (18 female, M = 20.25 years, SD=1.48, range 18-24 years, 2 left-handed according to self-report, 13 right-dominant eye). All participants had normal or corrected-to-normal vision (contact lenses only) and received $20 for their participation. One participant had to be replaced due to technical difficulties.

### Vocal Responses in the Continuous Flash Suppression Task

In Experiment 2, participant gave vocal instead of manual responses to indicate target detection and location. To this end we used a voice key. Participants were instructed to respond with a ‘ba’ sound as soon as the small target was detected, followed by a location categorization with another ‘ba’ sound or no sound. For half of the participants a second ‘ba’ sound indicated above fixation and for the other half it indicated below fixation. We used ‘ba’ as this gave a consistent sharp consonant sound to trigger the voice key reliably.

Participants were 97.48% (SD=3.06) correct across trials; in 0.63% of trials there was no response, and data trimming as described for Experiment 1 led to the removal of a further 1.56% of the trials.

## Results Experiment 2

Vocal reaction times in Experiment 2 were generally slower compared to the manual reaction times in Experiment 1. This is in line with other studies comparing these response modalities (Eimer & Schlaghecken, 2001). As can be seen in Figure 2B, we found the fastest reaction times for the hand masks (hands mean RT= 1733 ms, SD=643 ms). We also found relatively fast reaction times again for the tool masks (tools mean RT=1863 ms, SD=855 ms; hammers mean RT=1788 ms, SD=781 ms). In this experiment, the reaction times in the large object conditions were also similar to the tool conditions (large objects mean RT=1813 ms, SD=598 ms; small objects mean RT=1919 ms, SD=606 ms; phone mean RT=2054 ms, SD=782 ms; car mean RT=2119 ms, SD=763 ms). Nevertheless, we found significant data-model correlations for the hands/tools inefficient mask model (Bonferroni corrected significance threshold of *p*=0.025, tau-a mean=0.212, *p*<0.001). These correlations were significantly higher than the high-level representational architecture model correlations (permutation test p-values for the difference: *p*<0.001), for which we did not find significant data-model correlations (Bonferroni corrected significance threshold of *p*=0.025, tau-a mean=−0.0003, *p*=0.99). Overall, even with a non-manual response modality, we found the fastest breakthrough times for the hand masks ruling out a simple manual response facilitation explanation. Furthermore, we found no significant correlations for the high-level representational architecture model and found that the hand/tool inefficient mask model yielded better data-model correlations.

### Results for local category-specific image characteristics (Experiments 1 and 2)

As we found the lowest CFS breakthrough times for hands and tools, it is possible that these categories contain the lowest edge content of our stimuli. To estimate the number of edges in each image, we used the Matlab edge detection function. We employed the widely used *Canny* method with the low and high thresholds 0.1 and 0.2 (Canny, 1986). We also analysed the edges employing the *prewitt* method (Prewitt, 1970) and found comparable results, suggesting that the pattern of results was not dependent on the particular choice of edge detector. We calculated the average percentage of edges for all images across each category. We used these edge-content mean values to create edge-content models as predictors for breakthrough times for different category masks (Figure 3).

Although hammers and tools had the lowest object coverage in the target area, most likely due to their elongated shape, this was not the case for hands. We obtained significant data-model correlations for object coverage models (considering comparisons for five models, Bonferroni corrected significance threshold of *p*=0.01; Experiment 1 %object entire image tau-a mean=0.085, *p*<0.001, %object target image tau-a mean=0.148, *p*<0.001; Experiment 2: %object entire image tau-a mean=0.065, *p*=0.003, %object target image tau-a mean=0.178, *p*<0.001). However, our analysis also revealed that the data-model correlations for the object coverage for the entire image were significantly smaller compared to the hand/tool inefficient mask model (Experiment 1: tau-a mean=0.191; Experiment 2: tau-a mean=0.212, permutation test p-values for the difference, considering all four paired comparisons for object coverage and edge content models Bonferroni corrected significance threshold of p=0.0125: Experiment 1: %object entire image *p*<0.001, %object target area *p*=0.023; Experiment 2: %object entire image *p*<0.001, %object target area *p*=0.076).

Hands had the least amount of edges both across the entire image and specifically in the target area. Tools and hammers also had a lower edge content compared to several other categories. The correlations between data and edge-content models for both experiments were significantly above zero (considering comparisons for five models Bonferroni corrected significance threshold of *p*=0.01; Experiment 1 %edge entire image tau-a mean=0.231, *p*<0.001, %edge target area =0.217, *p*<0.001; Experiment 2: %edge entire image tau-a mean=0.303, *p*=0.002, %edge target area tau-a mean=0.292, *p*<0.001). The edge-content model correlation coefficients (except for the edge-target area model in Experiment 1) were also significantly larger than for the hands/tools inefficient mask model (Experiment 1: tau-a mean=0.191; Experiment 2: tau-a mean=0.212, permutation test p-values for the difference, considering all four paired comparisons for object coverage and edge content models Bonferroni corrected significance threshold of *p*=0.0125: Experiment 1: %edge entire image *p*<0.001, %edge target area *p*=0.088; Experiment 2: %edge entire image *p*<0.001, %edge target area *p*<0.001). Thus, this exploratory analysis suggests that the category-specific edge content provides the best predictor for the data and could underlie the hand and tool specific effects.

To further investigate if category-specific image characteristics can account for the hand and tool specific effects, we also ran partial correlation analyses. These allowed us to test data-model correlations for the hands/tools inefficient mask model while controlling for other variables such as object coverage and edge content. For both Experiment 1 and Experiment 2, we found that the correlation coefficients were reduced (Experiment 1: tau-a mean=0.154, *p*<0.001; Experiment 2: tau-a mean=0.120, *p*<0.001) but still significant when taking all four variables (object coverage entire image and target area, edge content entire image and target area) into account. This was also the case when we ran partial-correlation analyses for all four variables separately (Bonferroni corrected significance threshold of p=0.0125, Experiment 1: %object entire image tau-a mean=0.264, *p*<0.001; %object target area tau-a mean=0.237, *p*<0.001; %edge entire image tau-a mean=0.141, *p*<0.001; %edge target area tau-a mean=0.169, *p*<0.001; Experiment 2: %object entire image tau-a mean=0.305, *p*<0.001; %object target area tau-a mean=0.248, *p*<0.001; %edge entire image tau-a mean=0.102, *p*<0.001; %edge target area tau-a mean=0.148, *p*<0.001). This analysis suggests, that category-specific image characteristics such as edge content cannot fully explain the hand and tool specific effects.

In Experiment 3 we tested whether the edge-content explanation could contribute to the results of Cohen et al. (2015) by using the same categories as they employed.

## Experiment 3

In Experiment 3, we asked whether we also find relative inefficiency of hand masks when including other object categories such as those employed by Cohen et al. (2015). This also allowed us to test the edge-content model on another set of categories. Furthermore, we could test if we can find a comparable CFS breakthrough time pattern as reported by Cohen et al. (2015) and investigate to what extent it might be related to the amount of category-specific edge content.

### Participants

We tested 20 new participants (18 female, M = 23.55 years, SD=4.57, range 18-36 years, all right-handed according to self-report, 10 right-dominant eye). All participants had normal or corrected-to-normal vision (contact lenses only) and received Euro 7.50 for their participation. This research was conducted in accordance with the ethical standards laid down in the 1964 Declaration of Helsinki and was approved by the Ethics Committee of the Faculty for Social and Behavioural Sciences, Friedrich Schiller University Jena. Written informed consent was obtained from all participants prior to the start of the experiment.

### Methods

In Experiment 3, we employed the same methods as in Experiment 1 with a few exceptions. First, the study was conducted in Germany and thus we translated the instructions and feedback into German. Stimuli were presented on a BENQ monitor with a refresh rate of 60 Hz and a screen resolution of 1680 x 1050. In addition to the hand category, we included face, body, building, car and chair category sets.

Participants were 98.66% (SD=0.91) correct across trials; in 0.20% of trials there was no response, and data trimming as described above led to the removal of 1.56% of the trials.

## Results Experiment 3

First, we present our behavioural data as in Cohen et al. (2015; Figure 4), including only the categories of faces, bodies, buildings, cars and chairs, but separating masks and targets as in our previous experiments. As can be seen in Figure 4A, there are general mask effects (car masks have overall longer breakthrough times compared to other masks) and also general target effects (face targets have overall shorter breakthrough times compared to other targets). This suggest that the data pattern is not symmetrical across different mask and target pair combinations (e.g. face mask and body target pairs longer breakthrough times compared to body mask and face target pairs). We then transformed our data so it had the same format as Cohen et al. (2015): We excluded within-category mask and target pairs and collapsed across mask and target pairs (e.g., face mask and body target averaged with body mask and face target) (Figure 4A). The general effects between different masks and targets are no longer obvious in the collapsed data but they may nevertheless underlie the pair rank order. This highlights the importance of showing mask /target pairs separately in a non-collapsed format (for example, using our mosaic plots).

The collapsed pairs can be seen in a bar graph (Figure 4B) as in Cohen et al. (2015, Figure 4B). Our data showed a similar range of breakthrough times (from 1724 to 2010 ms). They also show a similar pattern with faces/buildings, faces/bodies and faces/ chairs among the lowest and cars/bodies, cars/buildings and cars/chairs among the highest breakthrough times. In contrast, we found longer breakthrough times for bodies & buildings compared to Cohen et al. (2015). We correlated the rank order of our data with theirs and found a significant correlation (Spearman’s rho = 0.71, *p*=0.019).

We then derived a model based on the rank order of the data by Cohen et al. (2015) with faces/buildings the highest rank (1^st^ rank = fastest breakthrough times) and car/chair pairs the lowest rank (10^th^ rank = slowest breakthrough times) (see Figure 4C). We derived the corresponding edge content and object coverage models by collapsing values across mask and target pairs. Using a correlation analysis between the models and our data (considering comparisons for five models Bonferroni corrected significance threshold of *p*=0.01) we found significant correlations between our data and the Cohen et al. order model (tau-a mean=0.2, *p*< 0.001). We also found significant correlations for the edge-content models (for both tau-a mean =0.2, *p*<0.001), but not the object-coverage models (%object entire image tau-a mean=−0.044, *p*=0.435 %object target area tau-a mean=−0.053, *p*=0.350). The edge-content model correlation coefficients were numerically and statistically not different to the Cohen et al. (2015) order model correlations (permutation test p-values for the model differences, Bonferroni corrected significance threshold of *p*=0.025: Cohen et al. versus %edge entire image *p* = 0.9774, Cohen et al. versus %edge target area *p* = 0.9774). This suggests that both the specific category order (related to category neural similarity) and the edge content are able to account for the data pattern in Experiment 3.

We also ran partial correlation analyses. These allowed us to test data-model correlations for the Cohen et al. order model while controlling for other variables such as object coverage and edge content. We found that the correlation coefficient was reduced (tau-a mean=0.173, *p*=0.002) but still significant when taking all four variables (object coverage entire image and target area, edge content entire image and target area) into account. This was also the case when we ran partial-correlation analyses for all four variables separately (Bonferroni corrected significance threshold of p=0.0125, %object entire image tau-a mean=0.210, *p*<0.001; %object target area tau-a mean=0.204, *p*<0.001; %edge entire image tau-a mean=0.180, *p*<0.001; %edge target area tau-a mean=0.180, *p*<0.001). This analysis suggests that category-specific image characteristics such as edge content cannot fully explain the category-specific effects. Thus it remains plausible that additional high-level representational or other factors influence visual awareness in CFS.

Overall, we found a similar pattern to the behavioural findings of Cohen et al. (2015) and found that edge-content was a good predictor for this pattern. These category-specific low-level image characteristics could not completely account for the behavioural data pattern, suggesting high-level factors can also contribute to the limitations on visual awareness.

In a second analysis, we included the data for the hand category. In line with Experiments 1 and 2, we found that hands were the most inefficient masks (shortest mean breakthrough times, RT mean for hands=1548 ms, SD = 473 ms; mean mask breakthrough times for other categories: faces mean RT= 1831, SD=473 ms; bodies mean RT=1798, SD=575 ms; buildings mean RT=1866, SD=531 ms; cars mean RT=2086, SD=746 ms; chairs mean RT=1815, SD=508 ms) and numerically had the lowest edge content in the context of the object categories employed by Cohen et al. (2015). In our first analysis (Figure 5), we compared the full data set (including hands) to hands-inefficient, edge-content and object-coverage models. We found the largest data-model correlations for the edge-content models (considering comparisons for five models Bonferroni corrected significance threshold of *p*=0.01; %edge target area tau-a mean=0.240, *p*<0.001; %edge entire image tau-a mean=0.226, *p*<0.001; %edge hand inefficient mask model tau-a mean =0.148, *p*<0.001; %object entire image tau-a mean=−0.002, *p*=0.945, %object target area tau-a mean=0.079, *p*=0.002). These were significantly larger than the correlations for the hands-inefficient mask model (permutation test p-values for the model differences, Bonferroni corrected significance threshold of *p*=0.025: %edge entire image *p*<0.001, %edge target area *p*<0.001). Thus, in line with the previous experiments edge-content models were the best predictor for the mask category reaction time order.

As in previous analyses, we also investigated if category-specific image characteristics could account for hand-specific effects using a partial correlation analysis. Again, we found that the correlation coefficient was still significant (tau-a mean=0.134, *p*<0.001) when taking all four image-specific variables (object coverage entire image and target area, edge content entire image and target area) into account. This was also the case when we ran partial-correlation analyses for all four variables separately (Bonferroni corrected significance threshold of p=0.0125, %object entire image tau-a mean=0.294, *p*<0.001; %object target area tau-a mean=0.282, *p*<0.001; %edge entire image tau-a mean=0.176, *p*<0.001; %edge target area tau-a mean=0.165, *p*<0.001). In line with the previous analyses, this suggests that category-specific image characteristics cannot fully explain the hand-specific effects.

## Discussion

We conducted three CFS experiments to explore the contributing factors to suppression of visual awareness across different object categories. In particular, we investigated this in the context of hand and tool stimuli. These stimuli form a particularly useful test category for testing the hypothesis that overlapping neural representations result in greater competition (Cohen et al., 2015) because neural representations for hands and tools have been shown to overlap (Bracci et al., 2012). We predicted longer CFS breakthrough times for hand/tool pairs compared to hand/other object pairs due to their neural overlap. We also explored the contribution of lower-level factors such as image coverage and edges.

In contrast to the predictions from the high-level neural representational architecture hypothesis, in Experiment 1, we found that participants were generally faster at detecting targets when paired with either hand or tool CFS masks, compared to other object category masks. These data did not correlate with a model based on higher-level neural representational architecture, but instead seemed to indicate a model where hands and tools were inefficient masks relative to other categories. We then tested this inefficiency model against new data in Experiment 2 and found significant positive correlations. We also found support for hands to be relatively inefficient masks in the context of those stimuli used in a previous CFS experiment by Cohen et al. (2015), finding a similar pattern as their behavioural CFS findings (Experiment 3).

When investigating the underlying mechanisms for these effects in more detail, we verified that the relative inefficiency of hands and tools was not due to facilitation of manual responses: we replicated the effect using vocal responses (Experiment 2). Analysis of category-specific local image characteristics showed that edge content (i.e., average percentage of mask image that is edge) but not object coverage (i.e., average percentage of mask image that is covered by a visual object) was the best predictor for category differences in breakthrough times across all analyses. Hand stimuli had, on average, the lowest edge content and this might be the underlying factor as to why hands are relatively inefficient masks. Edge content was also a good predictor for the data analysed without hands and collapsed across category pairs as in Cohen et al. (2015). Thus, our findings show that edge content is a relevant source for limitation in the CFS task with stimuli for which other low-level factors have been carefully controlled (e.g., luminance, contrast and colour).

In previous work, increased spatial density was related to increased CFS breakthrough times but it was not possible to tease apart effects of edge content and general contrast energy (Drewes et al., 2018). Here we used images with matched global spatial frequency content, contrast and luminance (Willenbockel et al., 2010) and found that edge content significantly correlated with breakthrough times across different category sets. This demonstrates that increased edge-detection workload itself can be linked to CFS breakthrough times. Edge detection happens very early in the visual processing stream both at the subcortical level (lateral geniculate nucleus in the thalamus) and in early visual cortex (Hubel & Wiesel, 1962). Thus, the specific mechanism that links edge content and suppression times is likely related to a low-level visual mechanism. Overall, our findings provide novel evidence for a low-level perceptual bottleneck for visual awareness in the CFS task.

Our findings in Experiments 1 and 2 do not provide evidence for a high-level mechanism limiting visual awareness for competing objects. In Experiment 3, we found a similar CFS pattern as reported by Cohen et al. (2015) using the same category sets. These authors showed high correlations between this pattern and the neural category similarity in high-level visual cortex. However, in Experiments 1 and 2 we were not able to predict, based on known neural similarity, the CFS pattern for a different set of categories involving hand, tool and object stimuli. A limitation of our study was that we based our sample size on the study by Cohen et al. (2015) and not on a statistical power analysis. We developed a novel method to analyse category-specific CFS breakthrough times which involves correlating our CFS data pattern with several different models. To determine whether these correlations are statistically significant we used permutation tests. Power analyses for these permutation tests would require large simulation studies and computing times beyond a feasible timeframe. Thus, although with our methods we were able to detect certain effects, including a similar pattern as reported in Cohen et al. (2015), it is still possible that our study may have not had enough statistical power to detect smaller effects. We note, however, that the average correlation coefficient and effect size for the high-level representational architecture model correlation was very close to zero (Experiment 1 tau-a mean=−0.002, Experiment 2 tau-a mean=−0.0003), suggesting that any effect we may have missed is likely to be very small.

Instead of support for the neural representational architecture model, we found general mask effects that correlated with edge content across all experiments. Our new visualisation and correlation methods allowed us to detect and investigate such general mask effects, whereas previous work used averages across mask and target pairs (e.g. face mask and body target averaged with body mask and face target; Cohen et al., 2015), which means there is no potential to detect differences between specific category masks or targets.

Although the edge account had good explanatory power, there was additional variance not explained by the edge model, leaving room for additional contributing factors. Our partial correlation analyses testing if category-specific image characteristics could account for the Cohen et al. (2015) category pattern or for the hand /tool effects resulted in reduced, but still significant, data-model correlation coefficients when controlling for edge content. This suggests that there may be both low- and high-level object category-specific limits for visual awareness. These might include contributions from high-level neural representational architecture (although we were not able to find any support for this model using hand/tool categories here), category-specific edge content and also special mask effects for specific categories such as hand and tool stimuli. Alternatively, it may be the case that using category means for edge content is not ideal to estimate the influence of edge content on the reaction time data, as edge content could also vary substantially within and between trials. To fully test for high-level contributions to the limits on visual awareness, it would be necessary to control edge content across categories or model it for individual trials.

Our lack of evidence for the high-level representational architecture model in Experiments 1 and 2 could also be stimulus- and task-specific. The CFS paradigm involves fast presentations of large masks over a relatively long trial time. This would challenge processing capacity in early visual cortex, which could then result in large effects of edge content on reaction time measures (Drewes et al., 2018), especially when using stimuli with relative high or low edge-content such as hands and tools. This could, in turn, mask other potential effects due to the representational architecture. Evidence for a link between the high-level representational architecture and visual processing of object stimuli has also been reported in paradigms where the object stimuli were spatially distributed (visual search paradigm: Cohen et al., 2017; visual memory paradigm: Cohen et al., 2014). It is possible that these multiple object tasks tap mostly into selective attention mechanisms that are indeed influenced by the neural high-level spatial distribution of the processing of visual categories. However, it is important to investigate the role of edge content also for these tasks to rule out low-level perceptual explanations.

It is also important to note that we only indirectly tested the high-level representational architecture based on what is known about the overlap between hand and tool areas and previous paired-category findings by Cohen et al. (2015). We used existing neural data to derive predictions in line with the predictions of the high-level architecture theory. In the future, it will be important to test behaviour-brain correlations of category-pair specific breakthrough times and neural similarity, but the current findings are important in showing such investigations should take edge content into account, and consider mask and target effects separately.

In this study we also found out more about the perceptual factors that might play a role in interactions between body and environment. We found that hands were less efficient masks compared to other categories, and that this is at least partly driven by the relative edge-content in these stimuli. Similarly, our tool stimuli tended to be less efficient masks and had relatively few edges compared to other categories. This could be specific to our exemplars but it seems unlikely, as generally hands and tools tend to have fewer details compared to many other categories of stimuli; this could be investigated in future studies including other stimulus sets. If it is true, due to visual object statistics, a low-level perceptual bottleneck could indirectly facilitate perception in body and tool interactions. And *vice versa*, such a potential mechanism could be taken into account when designing new artificial extensions for our bodies to be used for interactions. For example, artificial extensions should minimize any visible edges and details that are functionally not relevant.

Overall, our findings using hand stimuli as well as a range of other categories show that category-specific edge content influences the limits for visual awareness in the CFS paradigm with stimuli for which other low-level factors have been controlled (e.g., luminance, contrast, colour). In addition to these low-level limits, there may also be object category-specific high-level influences on visual awareness such as the neural representational architecture in higher visual cortex and hand- and tool-specific effects, but we need further research taking edge content into account to fully tease potential low- and high-level effects apart.

## Acknowledgements

RZ is supported by a Discovery Early Career Research Award from the Australian Research Council (ARC; DE140100499). ANR has research funding from the ARC (DP12102835 and DP170101840). We thank Marlene Wessels for assistance programming and data collection for Experiment 2. We thank Susan Wardle for helpful discussions and Serje Robidoux for statistical advice.

To access our stimuli, data, presentation and analyses scripts please use this Open Science Framework (OSF) link: https://bit.ly/2Krq7Ab.

## References

Almeida, J., Mahon, B. Z., Nakayama, K., & Caramazza, A. (2008). Unconscious processing dissociates long categorical lines. Proceedings of the National Academy of Sciences of the United States of America, 105(39), 15214–15218.

Bracci, S., Cavina-Pratesi, C., Ietswaart, M., Caramazza, A., & Peelen, M. V. (2012). Closely overlapping responses to tools and hands in left lateral occipitotemporal cortex. Journal of Neurophysiology, 107(5), 1443–1456. doi:10.1152/jn.00619.2011

Brainard, D. H. (1997). The Psychophysics Toolbox. Spatial Vision, 10(4), 433–436. doi:10.1163/156856897X00357

Brodeur, M. B., Dionne-Dostie, E., Montreuil, T., & Lepage, M. (2010). The Bank of Standardized Stimuli (BOSS), a new set of 480 normative photos of objects to be used as visual stimuli in cognitive research. PloS One, 5(5), e10773. doi:10.1371/journal.pone.0010773

Canny, J. (1986). A computational approach to edge detection. IEEE Transactions on Pattern Analysis and Machine Intelligence, 8, 679–698.

Cohen, M. A., Alvarez, G. A., Nakayama, K., & Konkle, T. (2017). Visual search for object categories is predicted by the representational architecture of high-level visual cortex. Journal of Neurophysiology, 117(1), 388–402. doi:10.1152/jn.00569.2016

Cohen, M. A., Dennett, D. C., & Kanwisher, N. (2016). What is the Bandwidth of Perceptual Experience? Trends in Cognitive Sciences, 20(5), 324–335. doi:10.1016/j.tics.2016.03.006

Cohen, M. A., Konkle, T., Rhee, J. Y., Nakayama, K., & Alvarez, G. A. (2014). Processing multiple visual objects is limited by overlap in neural channels. Proceedings of the National Academy of Sciences of the United States of America, 111(24), 8955–8960. doi:10.1073/pnas.1317860111

Cohen, M. A., Nakayama, K., Konkle, T., Stantic, M., & Alvarez, G. A. (2015). Visual Awareness Is Limited by the Representational Architecture of the Visual System. Journal of Cognitive Neuroscience, 27(11), 2240–2252. doi:10.1162/jocn_a_00855

Desimone, R., & Duncan, J. (1995). Neural mechanisms of selective visual attention. Annual Review of Neuroscience, 18, 193–222.

Drewes, J., Zhu, W., & Melcher, D. (2018). The edge of awareness: Mask spatial density, but not color, determines optimal temporal frequency for continuous flash suppression. Journal of Vision, 18(1), 12. doi:10.1167/18.1.12

Eimer, M., & Schlaghecken, F. (2001). Response facilitation and inhibition in manual, vocal, and oculomotor performance: evidence for a modality-unspecific mechanism. Journal of Motor Behavior, 33(1), 16–26. doi:10.1080/00222890109601899

Hesselmann, G., Darcy, N., Ludwig, K., & Sterzer, P. (2016). Priming in a shape task but not in a category task under continuous flash suppression. Journal of Vision, 16(3), 17. doi:10.1167/16.3.17

Hubel, D. H., & Wiesel, T. N. (1962). Receptive fields, binocular interaction and functional architecture in the cat’s visual cortex. Journal of Physiology, 160, 106–154.

Kim, C. Y., & Blake, R. (2005). Psychophysical magic: rendering the visible ‘invisible’. Trends in Cognitive Sciences, 9(8), 381–388. doi:10.1016/j.tics.2005.06.012

Kim, S. (2015). ppcor: An R Package for a Fast Calculation to Semi-partial Correlation Coefficients. Commun Stat Appl Methods, 22(6), 665–674. doi:10.5351/CSAM.2015.22.6.665

Konkle, T., & Oliva, A. (2012). A real-world size organization of object responses in occipitotemporal cortex. Neuron, 74(6), 1114–1124. doi:10.1016/j.neuron.2012.04.036

Kriegeskorte, N., Mur, M., Ruff, D. A., Kiani, R., Bodurka, J., Esteky, H., … Bandettini, P. A. (2008). Matching categorical object representations in inferior temporal cortex of man and monkey. Neuron, 60(6), 1126–1141. doi:10.1016/j.neuron.2008.10.043

Longo, M. R., & Haggard, P. (2009). Sense of agency primes manual motor responses. Perception, 38(1), 69–78.

Ludwig, K., Kathmann, N., Sterzer, P., & Hesselmann, G. (2015). Investigating category-and shape-selective neural processing in ventral and dorsal visual stream under interocular suppression. Human Brain Mapping, 36(1), 137–149. doi:10.1002/hbm.22618

Miles, W. R. (1930). Ocular dominance in human adults. Journal of General Psychology, 3, 412–430.

Nili, H., Wingfield, C., Walther, A., Su, L., Marslen-Wilson, W., & Kriegeskorte, N. (2014). A toolbox for representational similarity analysis. PLoS Computational Biology, 10(4), e1003553. doi:10.1371/journal.pcbi.1003553

Pelli, D. G. (1997). The VideoToolbox software for visual psychophysics: transforming numbers into movies. Spatial Vision, 10(4), 437–442.

Phipson, B., & Smyth, G. K. (2010). Permutation P-values should never be zero: calculating exact P-values when permutations are randomly drawn. Statistical Applications in Genetics and Molecular Biology, 9, Article39. doi:10.2202/1544-6115.1585

Prewitt, J. M. S. (1970). Object enhancement and extraction. In B. Lipkin & A. Rosenfels (Eds.), Picture Processing and Psychopictorics (pp. 75–149). New York: Academic.

Rosch, E., Mervis, C. B., Gray, W. D., Johnson, D. M., & Boyes-Braem, P. (1976). Basic objects in natural categories. Cognitive Psychology, 8, 382–439.

Stein, T., Sterzer, P., & Peelen, M. V. (2012). Privileged detection of conspecifics: evidence from inversion effects during continuous flash suppression. Cognition, 125(1), 64–79. doi:10.1016/j.cognition.2012.06.005

Tong, F., Meng, M., & Blake, R. (2006). Neural bases of binocular rivalry. Trends in Cognitive Sciences, 10(11), 502–511. doi:10.1016/j.tics.2006.09.003

Treisman, A. M., & Gelade, G. (1980). A feature-integration theory of attention. Cognitive Psychology, 12, 97–136.

Tsuchiya, N., & Koch, C. (2005). Continuous flash suppression reduces negative afterimages. Nature Neuroscience, 8(8), 1096–1101. doi:10.1038/nn1500

Tsuchiya, N., Koch, C., Gilroy, L. A., & Blake, R. (2006). Depth of interocular suppression associated with continuous flash suppression, flash suppression, and binocular rivalry. Journal of Vision, 6(10), 1068–1078. doi:10.1167/6.10.6

Willenbockel, V., Sadr, J., Fiset, D., Horne, G. O., Gosselin, F., & Tanaka, J. W. (2010). Controlling low-level image properties: the SHINE toolbox. Behavior Research Methods, 42(3), 671–684. doi:10.3758/BRM.42.3.671

Wolfe, J. M., Cave, K. R., & Franzel, S. L. (1989). Guided search: an alternative to the feature integration model for visual search. Journal of Experimental Psychology: Human Perception and Performance, 15(3), 419–433.

Zhu, W., Drewes, J., & Melcher, D. (2016). Time for Awareness: The Influence of Temporal Properties of the Mask on Continuous Flash Suppression Effectiveness. PloS One, 11(7), e0159206. doi:10.1371/journal.pone.0159206

